# Spatial lipidomics of fresh-frozen spines

**DOI:** 10.1101/2023.08.23.554488

**Authors:** Kayle J. Bender, Yongheng Wang, Chuo Ying Zhai, Zoe Saenz, Aijun Wang, Elizabeth K. Neumann

**Author notes:** **Corresponding Author** Elizabeth K. Neumann - Department of Chemistry, University of California, Davis, One Shields Avenue, Davis, CA 95616, United States.

## Abstract

Technologies assessing the lipidomics, genomics, epigenomics, transcriptomics, and proteomics of tissue samples at single-cell resolution have deepened our understanding of physiology and pathophysiology at an unprecedented level of detail. However, the study of single-cell spatial metabolomics in undecalcified bones faces several significant challenges, such as the fragility of bone which often requires decalcification or fixation leading to the degradation or removal of lipids and other molecules and. As such, we describe a method for performing mass spectrometry imaging on undecalcified spine that is compatible with other spatial omics measurements. In brief, we use fresh-freeze rat spines and a system of carboxyl methylcellulose embedding, cryofilm, and polytetrafluoroethylene rollers to maintain tissue integrity, while avoiding signal loss from variations in laser focus and artifacts from traditional tissue processing. This reveals various tissue types and lipidomic profiles of spinal regions at 10 μm spatial resolutions using matrix-assisted laser desorption/ionization mass spectrometry imaging. We expect this method to be adapted and applied to the analysis of spinal cord, shedding light on the mechanistic aspects of cellular heterogeneity, development, and disease pathogenesis underlying different bone-related conditions and diseases. This study furthers the methodology for high spatial metabolomics of spines, as well as adds to the collective efforts to achieve a holistic understanding of diseases via single-cell spatial multi-omics.

---

The spinal column, or vertebral column, is important in both structure and function in many organisms, including humans^1^. The main components of the spinal column are the spinal cord, vertebrae, intervertebral discs^2^, muscle, and blood vessels, which combine to enable communication between the brain and the body for sensory and motor function^3^. There are several layers to the spinal cord, including the grey matter in the inner part of the spinal cord and white matter in the outer part^4^. Grey matter of the spinal cord contains nuclei involved in sensory and motor functions and the white matter contains the tracts of axons, known as the ascending tracts and descending tracts^5^. Information is in the spinal cord is carried either to the brain via ascending tracts or to the motor neurons via descending tracts.^5^ The vertebrae of the spinal column are bones that surround and protect the spinal cord and roots^6^. There are many diseases or injuries that affect the spinal column, often resulting in changes or loss in function(s) due to its importance and complexity^7,8^. These dysfunctions vary in severity and are broadly categorized as mechanical, neuropathic, or medical^8^ and include spina bifida^9^, scoliosis^10^, rheumatoid arthritis^11^, osteoporosis^12^, and spinal cord injury (SCI)^13^. SCI is characterized by neurological dysfunction and may or may not include damage to the rest of the spinal column^14^. Depending on the nature of the trauma and the location within the spine at which the trauma occurred, SCI due to trauma to the spinal column differs in severity^15^. Whether the SCI is chronic or not, there are many associated conditions^14^. Notably, even acute SCI can cause dysregulation of cardiovascular function due to the loss of sympathetic control^14^. Because of the importance and possible dysfunctions of the spine, it is necessary to have ways to study the spine.

The spine may be imaged in humans for diagnostic purposes, which may be accomplished via several modalities of various sensitivities and resolution^16^. The most common is x-ray radiography^17^, which quickly (sec-min) provides information for a m^2^ sized areas compared to other related modalities^18^ and provides radiographic information about bone structures in a non-static context such as mobility^19^ or bending radiography^20,21^. Computed tomography (CT) involves reconstructing the internal structures of the body from a collection of X-ray data taken from hundreds or more angles^22,23–25^. Both X-ray radiography and CT share the risks of high radiation exposure, especially for higher-resolution images^26^. Another form of imaging is nuclear magnetic resonance imaging (MRI), which defines soft tissues and neural elements^27^ and can non-invasively create an image without applying radiation^28,29^ through the use of dyes^30^. The above methods of imaging can be performed in living organisms, but lack discrete molecular resolution, which is critical in understanding healthy and diseased (dys)function. Some methods may be more suited for discrete molecular information, albeit at the cost of imaging are, include Raman spectroscopy^31^, infrared spectroscopy^32^, and mass spectrometry imaging^33^, which use spectral information at exact spatial locations to create an image with varying levels of specificity. To determine discrete metabolomic profiles of critical spinal cord regions, we use matrix-assisted laser desorption/ionization mass spectrometry imaging (MALDI MSI), which is a label-free imaging method that uses a laser to desorb/ionize molecules that have been co-crystallized in a small organic matrix^34^. Because this ionization method relies on the use of a laser, most of the ionized molecules come from the sample surface^34^. Because of the sensitivity and power of MSI, it can provide orthogonal information that could not be acquired via MRI or CT alone, such as the spatial location of lipids^35^, glycans^36^, peptides^37,38^, proteins^39–41^, and metabolites^42–44^ in tissues. Previously, MALDI MSI been applied to diseases^45–47^, dysfunction^48^, development^49^, trauma^50,51^, drug discovery and development^52–54^, etc. by using animal^46–55^ and human^45,55,56^ models, demonstrating its utility and importance.

Prior work on fresh-frozen tissue was performed by separating the spinal cord from the surrounding bone, then using desorption electrospray ionization (DESI) MSI^57–59^, secondary ion mass spectrometry (SIMS) MSI^60,61^, or MALDI MSI^60,62^. As such, we have aimed to image non-decalcified, fresh frozen spinal cord while it is still surrounded by the corresponding bone and muscle to best preserve and visualize its native metabolic features. In brief, the MALDI MSI workflow requires successful tissue preparation for data acquisition, making methods for tissue preparation an important area of research in the field^63,64^. Sectioning the tissue to a thickness of approximately 8-12 μm^65^, with thinner sections resulting in a better signal-to-noise ratio,^66^ is a major requirement of MALDI MSI. For this reason, a method for sectioning the spinal column without disturbing its native chemical architecture is necessary. Commonly, the spinal column is decalcified before sectioning to soften the bone, but decalcification interferes with the endogenous metabolites^67^. Methods for sectioning bone have been previously accomplished^68–72^, and we aim to develop subsequent methods to apply this approach to the spinal column. The spinal column is difficult to section to an 8-12 μm thickness because there are multiple tissue types of different densities and hardness being cut at a single time. Often, the sectioned spinal column does not maintain spatial integrity if cut and manipulated as loose tissue. Here, we demonstrate a method of performing MALDI MSI on fresh frozen, unfixed, undecalcified spinal columns at 10 μm spatial resolution. To do this, we use a cryotape-based method for sectioning the spinal column, including the spinal cord with surrounding bone and soft tissues intact. We compare the sectioning results with and without our approach to demonstrate the benefit of our approach with the goal of maintaining spatial and structural integrity of the spinal column during tissue preparation. This method is developed as a subsequent method of bone sectioning for MALDI MSI^68,72^ to be applied to the spinal column.

## EXPERIMENTAL SECTION

### Chemicals

Chemicals were purchased by Thermo Fisher and used without further purification unless otherwise specified.

### Tissue Preparation

All procedures were approved by the University of California Davis Institutional Animal Care and Use Committee and performed in accordance with the guidelines and policies of the Animal Center. Spines were harvested from Sprague Dawley female rats, aged 3 months (89 days) and euthanized with an overdose of carbon dioxide, with the strain code 400 sourced from Charles River. The spines were frozen using an isopentane-dry ice slurry and stored at -80°C. When specified, spines were embedded and frozen in carboxymethyl cellulose (CMC; EMD Millipore Corp., Burlington, MA). The tissue was sectioned to 10 μm using Cryofilm type 2C(9) tape (SECTION-LAB Co. Ltd., Yokohama Kanagawa, Japan) and a cryostat (Leica Biosystems, Wetzlar, Germany) while at -20 °C. The tissue in CMC was held in the cryostat using an optimal cutting temperature compound (Fisher HealthCare, Houston, TX). After sectioning, the tape with tissue was adhered to a slide prepared with copper tape (SECTION-LAB Co. Ltd.) and ZIG 2-way glue (Kuretake Co., Ltd., Nara, Japan) before being heat-fixed at 37 °C. Tissue was used immediately after sectioning.

### Staining

H&E staining (Abcam plc., Cambridge, UK) was performed at ∼20 °C (room temperature) as previously described, with minor alterations to the previous procedure to avoid damage to the glue and tape^73^. In brief, the tissue was incubated in hematoxylin, rinsed in water, incubated in bluing reagent, rinsed, and incubated in Eosin for contrast. Tissue was then coversliped with glycerol wet mount medium (Rs’ Science) for microscopic imaging using an EVOS™ M7000 Imaging System (brightfield, 20x objective; Invitrogen, Waltham, MA) was used for microscopy imaging.

### Matrix Application

1,5-Diaminonaphthalene (DAN, Tokyo Chemical Industry Company Ltd., Tokyo, Japan) matrix (20mg/mL in tetrahydrofurane) was applied using an HTX M3+ sprayer (HTX Technologies, LLC, Chapel Hill, NC). Important parameters include: 40 °C nozzle, gas pressure was 15 nitrogen psi, 50 μL/ min solvent flow rate, and a 2 mm track spacing for 5 passes.

### MALDI Q-TOF MSI

A timsTOF fleX dual source MALDI mass spectrometer (Bruker Scientific, Billerica, MA) was used for the MALDI MSI experiments. Each experiment was performed in positive mode with a 10 μm^2^ or 30 μm^2^ raster width as described in subsequent figure captions. A detailed description of the method is available in the supplemental information section (SI table S1). In brief, 150 shots were fired in a single burst, the global attenuator value was set to 0%, the local laser power was set to 88%, and the *m/z* range acquired was *m/z* 50 to 2000. The laser was manually focused using Galvano offset, moving the laser in the X and Y direction by up to 80 μm in each to compensate for the height of the tape.

### DataAnalysis

A. n internal, quadratic calibration of *m/z* values was performed using common lipids ([PC(26:0)+H], ([PC(32:0)+H], ([PC(32:0)+Na], ([PE(36:1)+H], ([PC(34:2)+H], ([PC(36:1)+Na]) within DataAnalysis (Bruker Scientific). Putative lipid identification was performed using the LIPID MAPS database^74^. All MALDI MSI ion images were created using SCiLS Lab Version 2023 (Bruker Scientific) with total ion count normalization. Average spectra were created by manually segmenting different regions using SCiLS Lab.

## RESULTS AND DISCUSSION

### Sectioning Workflow – Tape Method

Our goal was to develop a method for performing MALDI MSI on fresh frozen, unfixed, undecalcified spinal column (**Figure 1**). Ultimately, any tissue distortion or degradation will prevent high quality and reproducible spatial analysis by MALDI MSI or other imaging modalities. First, we embed and froze the undecalcified fresh-frozen spinal column in 2.6% CMC at -80 °C to preserve the chemical architecture of the tissue and protect the edges during sectioning. Spinal column is complex and contains diverse components which will not remain intact as a cohesive section once sectioned from the tissue block, but by adding tape, the section is reinforced such that a single, cohesive piece can be maneuvered. This reinforced spinal section can be manipulated similarly to other tissues, such as brain or kidney, enabling 10 μm sectioning. We apply conductive copper tape across the entire ITO coated glass slide and press the copper tape uniformly onto the slide using a razor blade. A thin and uniform layer of ZIG 2-way glue is added across the slide for adhering the tape and tissue. The addition of excess glue or air bubbles prevent laser focusing during MALDI MSI analysis. Additionally, if the glue is not dry, the glue layer often becomes uneven perhaps from the PTFE roller or other physical interactions. The slide is then cooled for at least 30 seconds to prevent heat fixation prior to tape adherence. One adhered, we gently use a PTFE roller to remove air bubbles, facilitating a consistent and flat surface for MALDI MSI. Finally, the slide is coated in DAN matrix and MALDI MSI is performed. Although we use DAN as a MALDI matrix, other matrices can be used with this method with minor adjustments to sprayer methods

**Figure 1.**
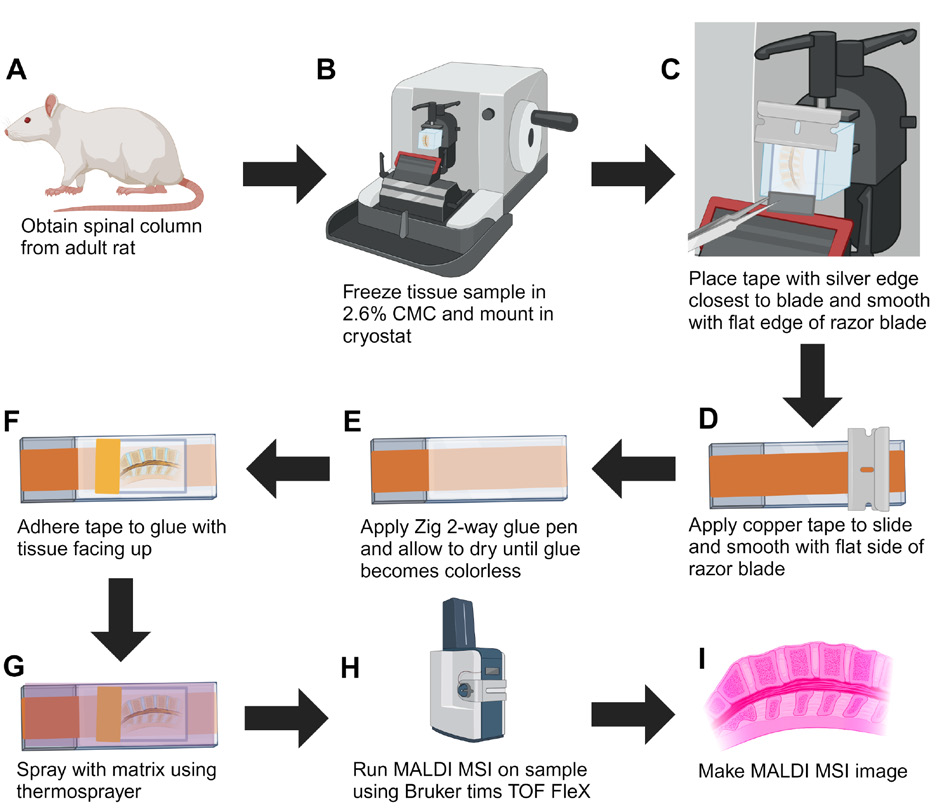
Workflow for preparing undecalcified fresh-frozen spinal column for MALDI MSI. (A) Lumbar tissue is excised from adult rats. (B) Tissue embedded in 2.6% CMC for enhanced cryosectioning. (C) The silver edge of the tape is placed over the edge of the blade. (D) A conductive copper tape is applied to the slide and adhered using a razor blade. (E) ZIG 2 way glue is applied on top the copper tape and allowed to dry until colorless. (F) The tape is adhered to the glue with the tissue on the side opposite from the glue. (G) Matrix is applied via an automated sprayer (HTX M3+) to enhance ion generation and desorption (H) MALDI MSI is performed for spatial lipidomics. (I) Data analysis is performed using a variety of software packages to create lipid profiles of major spinal regions. Figure was made using Biorender.

The above procedure has several variables that were tested and optimized to determine their effects on tissue integrity, using H&E staining as a metric. Here, we compare the sections with and without the use of a cryotape as well as with and without CMC embedding to assess the effects of each step. H&E staining shows the severity of vertebrae cracking and tissue rearrangement or destruction during sectioning. Overall, the spinal column sectioned using cryotape resulted in less bone cracking and tissue degradation than the equivalent sections, sectioned without cryotape (Figure 2). Additionally, embedding in CMC resulted in fewer cracks within the vertebrae when compared to non-embedded tissue, albeit this effect was not as significant as seen with use of the cryotape. Moreover, CMC serves a secondary benefit as the tape often gets caught on the blade when CMC is not present (Figure B.3). Although the cryotape has reduced the cracking and tissue degradation, there are some cracks that can be addressed in future experiments. Largely, the tissue remains intact as is evident in the clean boundaries shown by the H&E staining (Figure 2).

**Figure 2.**
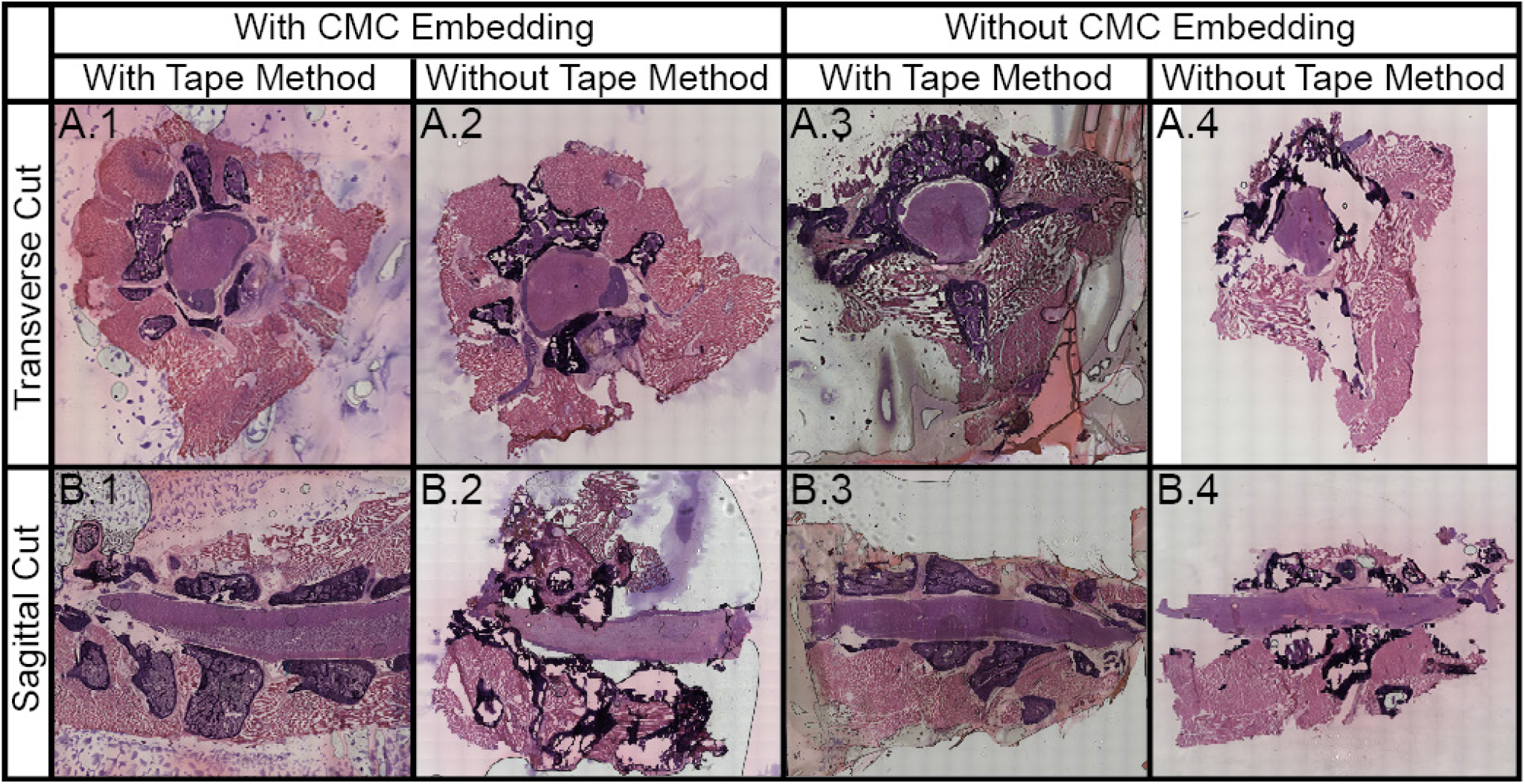
H&E stained section of undecalcified, unfixed, fresh-frozen spinal column under different conditions for both transverse and sagittal sections showing bone, bone marrow, muscle, nerves, grey matter, and white matter (SI Figure 1). In brief, we assessed the effects of CMC embedding and use of cryotape for both transverse (A) and sagittal cuts (B). In general, we found that CMC embedding and the use of the cryotape preserved the tissue adequately (left column). Cryotape was required for preserving the vertebra for both transverse (A.1 and A.3) and sagittal (B.1 and B.3) sections, while CMC embedding preserved the muscle tissue surrounding the spinal cord for both transverse (A.2 and A.4) and sagittal (B.2 and B.4) sections.

Spinal column sections without cryotape resulted tissue rearrangement and loss due to the fragility of the undecalcified bone and the innate separation between various tissue types throughout the spinal column. Various areas within the spinal column are not directly connected, so they move independently of each other. The tape directly adheres to the tissue, maintaining the integrity and spatial arrangement of the spinal cord. Additionally, the tape prevents the tissue from curling, which contributes to bone cracking and spatial rearrangement of the tissue, since it must be uncurled during slide adherence. The tissue blocks embedded in CMC allowed for a flat surface to adhere the entire piece, while the unembedded tissue has loose tape edges around the tissue that often caught on the blade during sectioning, resulting in additional artifacts. As such, the use of cryotape and CMC embedding preserves the most tissue content. Although other embedding media were not directly tested, they should be compatible with the use of cryofilm.

The tape is thin and flexible, making it difficult to adhere to the tape uniformly and resulting in the presence of air bubbles that will negatively affect MALDI MSI. To address this, we tried both pressing the tape to the glue with forceps and using a PTFE roller. In brief, the forceps were used to press the tape to the glue without directly touching the tissue, as direct contact by forceps would result in tissue damage. However, this resulted in air bubbles from inadequate tape adherence. Using a PTFE roller, however, resulted in fewer air bubbles, likely because a more uniform pressure could be applied to a larger area at a single time (Figure 3). Like other artifacts, air bubbles can be problematic for high spatial resolution MALDI MSI because any tissue above an air bubble is at a different height than the surrounding tissue, and thus, a different laser focus. This results in areas where the peak intensities are significantly lower or altogether absent (Figure 3D). We found that the PTFE roller must be used before heat fixation because the warm tissue will smear (Figure 3C), unlike when the tissue is still at -20 °C (Figure 3B). Using a PTFE roller successfully removes air bubbles with minimal damage to the tissue because the PTFE roller acts as a flat surface to apply consistent pressure across the tape and PTFE does not stick to tissue.

**Figure 3.**
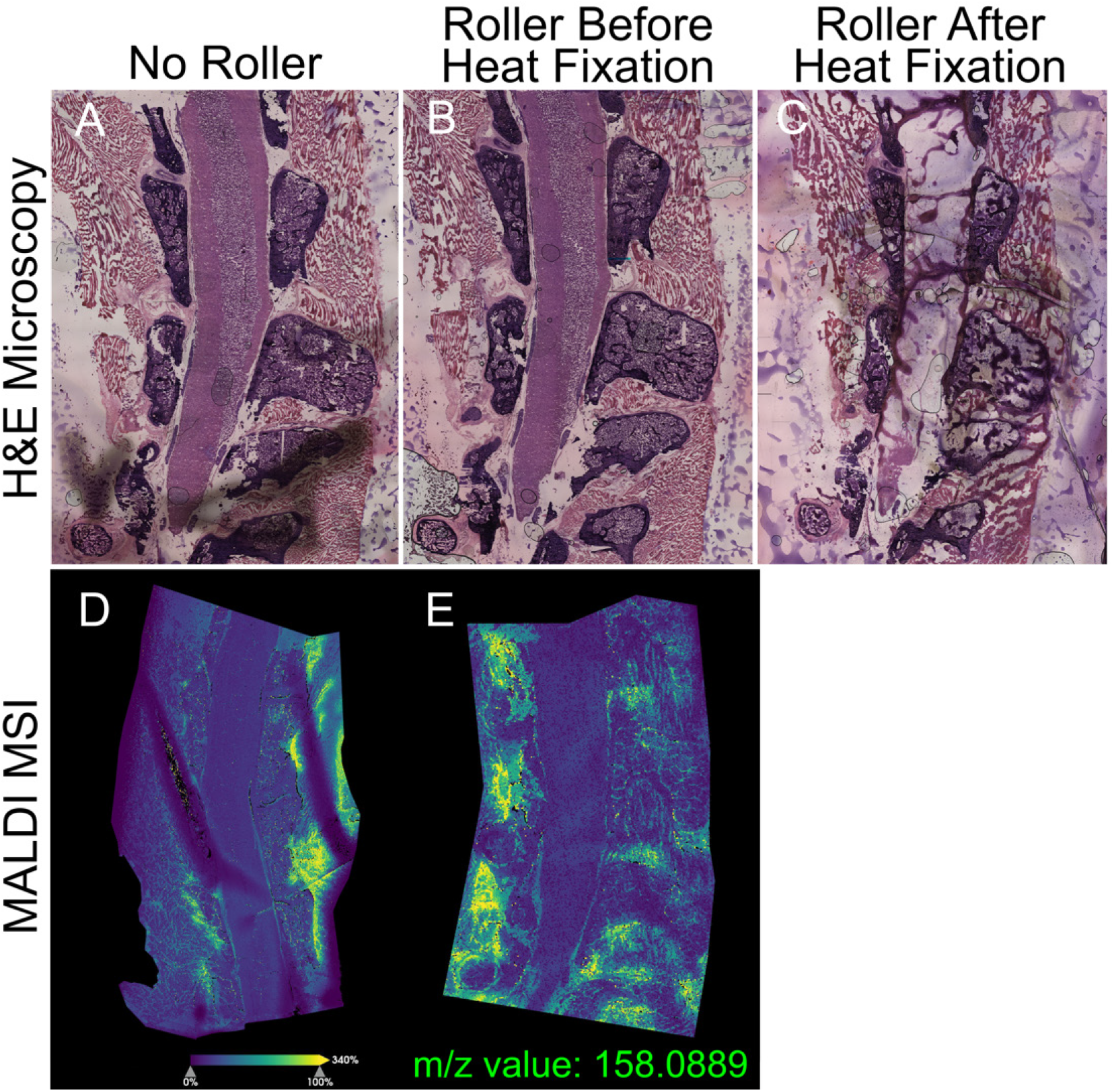
Use of a PTFE roller resulted in fewer air bubbles and, thus, fewer imaging artifacts. (A) H&E stain of a sagittal, undecalcified fresh-frozen spinal column embedded in CMC adhered to the slide using forceps without direct tissue contact. The presence of air bubbles between tape and glue is indicated by dark gray shadows. (B) An H&E stained tissue section adhered using a PTFE roller prior to heat fixation. (C) An H&E stained tissue section adhered using a PTFE roller after heat fixation, resulting in significant tissue degradation. (D) Example ion image of a sagittal spinal cord that was adhered using forceps as opposed to a roller (E).

On average, we detected 54 lipids in positive ion mode, with a majority being phosphatidylcholines (PC; SI Table S2), which is typical of positive ion mode. Although not performed, this sampling method is compatible with negative ion mode. The average mass spectrum for each tissue type shows a difference in ion composition (Figure 4). For instance, the bone marrow has an abundance of [PC(O-34:1)+H]^+^, [PC(O-34:1)+Na]^+^, [PC(34:1)+K]^+^ lipids, while the muscle is characterized by [PC(34:2)+K]^+^. Lipids play vital roles in a variety of cellular structures and processes because they are indispensable elements of myelin, cellular membranes, and vesicles that enable intracellular trafficking. In addition, lipids are involved in metabolic processes for energy production and are implicated in inflammations and bone-related diseases^75^. Largely, we are detecting membrane lipids, such as sphingomyelin (SM) and phosphatidylcholine (PC) lipids.^76^ Differences in lipid content may be a result of different cell types or cell shapes within these regions.^77–80^ For these reasons, we believe it is important to study the spinal lipidome. While there are discrete differences between tissue regions, the reproducibility of these profiles is notable (SI Figure S1). To demonstrate the localization of these lipids, we picked the top discriminators of each region (Figure 5).

**Figure 4.**
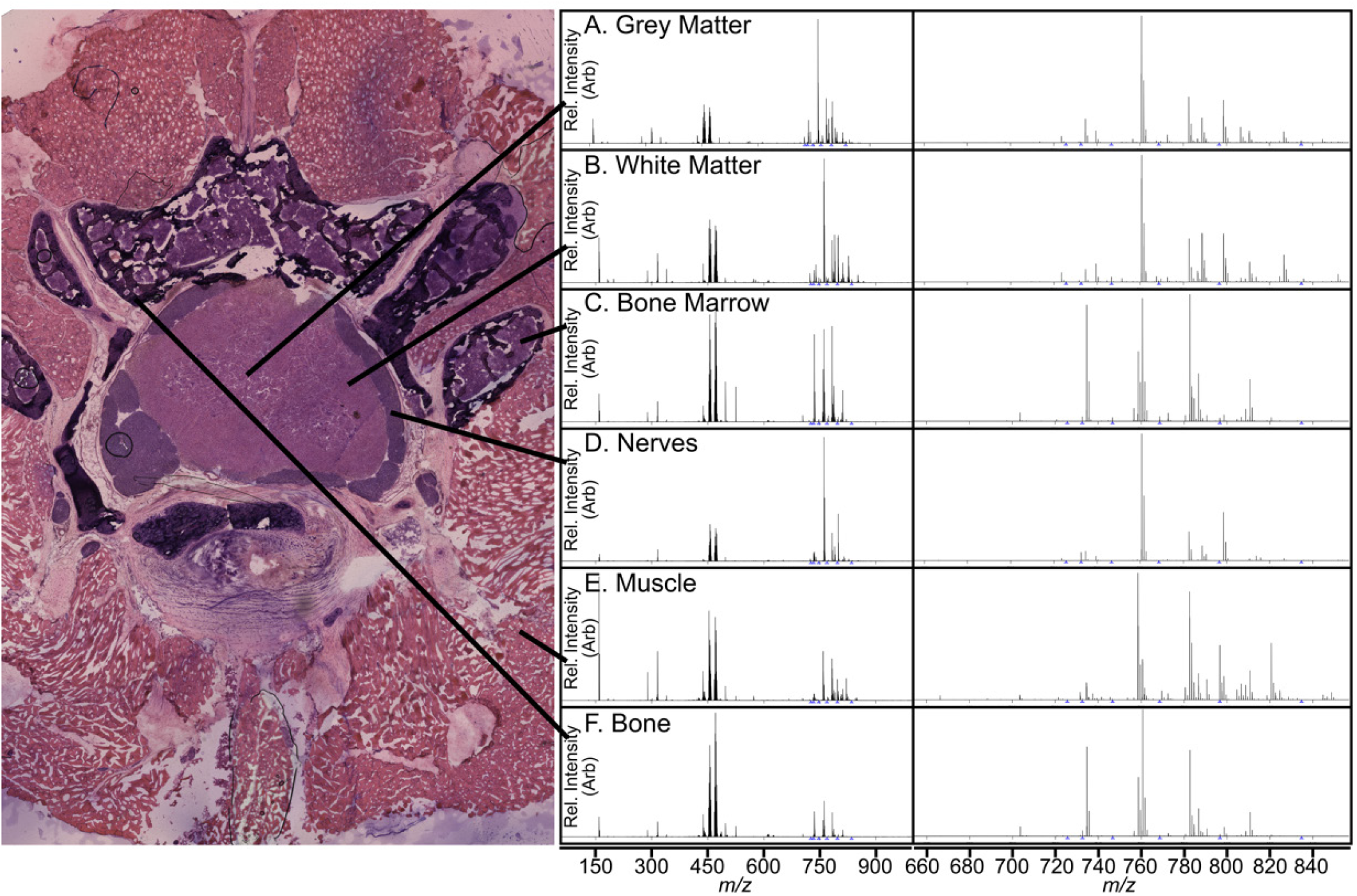
Average mass spectrum for each region within the spinal column (n = 3) with lines pointing to the respective region. The averaged mass spectrum is shown for each region (left) and enlarged spectra covering the lipid region (right).

**Figure 5.**
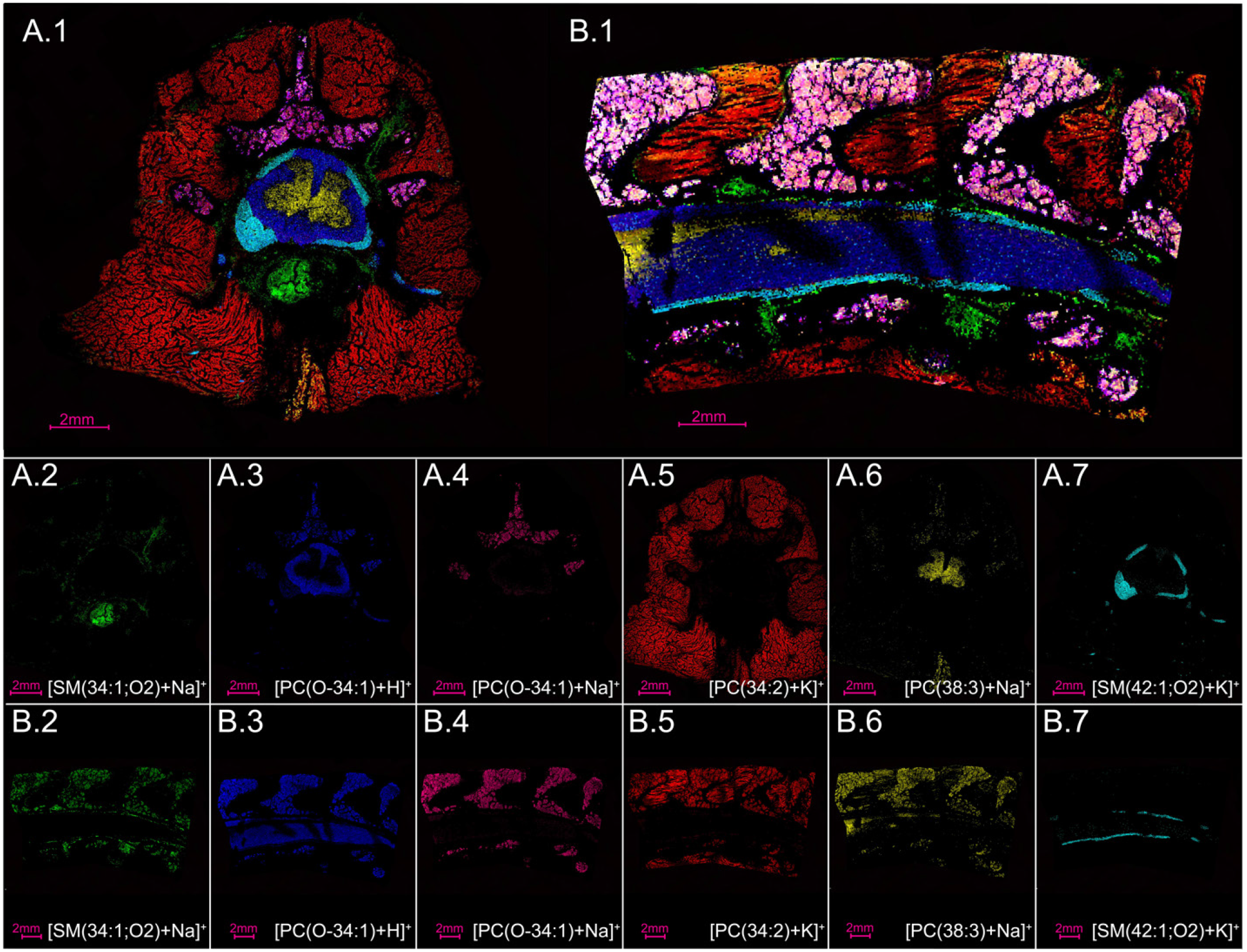
The major components of the spinal column can be visualized using MALDI MSI. such as intervertebral discs and blood vessels (A.2, B.2, *m/z* 725.5557, lipid [SM(34:1;O2)+Na]^+^), white matter (A.3, B.3, *m/z* 746.6041, lipid [PC(O-34:1)+H]^+^), bone marrow (A.4, B.4, *m/z* 768.5857, lipid [PC(O-34:1)+Na]^+^), muscle (A.5, B.5, *m/z* 796.5243, lipid [PC(34:2)+K]^+^), grey matter (A.6, B.6, *m/z* 834.5967, lipid [PC(38:3)+Na]^+^), nerves (A.7, B.7, *m/z* 853.6529, lipid [SM(42:1;O2)+K]^+^).

The sagittal cut results in a larger section than the transverse cut (Figure 5), making it more difficult to uniformly apply to the slide. This resulted in some taller regions of tape where the laser was less focused than it was during data collection of surrounding areas on the tissue (Figure 5B.1). Note, the sections in Figure 3E and and Figure 5B.1 are the same tissue slide and data set, with different ions and colors. The ions chosen for Figure 5 represent sphingomyelin (SM) and phosphatidylcholine lipids (PC), which are the two most common lipid classes in the outer cellular membrane.^76^ SM lipids are abundant in myelin sheaths, which electrically insulate nerve cell axons.^81^ [SM(34:1;O2)+Na]^+^ (green, Figure 5) is a sphingomyelin lipid that localizes to the intervertebral disc and blood vessels in both the transverse and sagittal cuts. Interesting, this lipid has been known to accumulate in the glomeruli of diabetic and high-fat diet fed mice,^82^ perhaps indicating that it may have a functional role in wildtype spinal cord, but a dysregulation or protective function in other organs. Another sphingomyelin lipid, [SM(42:1;O2)+K]^+^, localizes to nerves (teal, Figure 5), and could be a necessary component in neuronal membranes^83^ . SM lipids are, indeed, relevant to human health,^84^ demonstrated by their role in diabetes,^82,85,86^ coronary artery disease,^87,88^ Parkinson’s disease,^89^ cancer,^90–92^ and Niemann-Pick disease,^93^ and mapping them to specific functional regions may help decipher discrete SM functional roles. Additionally, [PC(38:3)+Na]^+^ localizes to the grey matter (yellow, Figure 5) and is also detected in lower abundance within the bone marrow and muscle. Moreover, [PC(O-34:1)+H]^+^ was detected within both the white matter and bone marrow (blue, Figure 5) and its sodiated form only within the bone marrow (pink, Figure 5), perhaps indicating a difference in salt content. [PC(34:2)+K]^+^) lipid (red, Figure 5) is localized mainly to the muscle, with some presence in the bone marrow. This is a significant lipid to detect because PC (34:2) is thought to be a biomarker for metabolic syndrome (MetS), which is related to an increased risk of diabetes and atherosclerotic cardiovascular disease (ASCVD).^94,95^

We aimed to assess spines from several rats at various points in the lumbar region to compare the ions in different tissue types. Three undecalcified fresh-frozen spinal columns from three different adult rats resulted in ion images that showed consistent localization of the same six ions based on the spatial location of the tissue types (Figure 6). The MALDI MSI layered ion images of the spinal columns cut in the transverse direction (Figure 6) show the same ions but result in images that appear slightly different overall. For instance, there appears to be more or less of any given tissue type and overall size in a section, each of which are from various places in the spine of three different rats (Figure 6). This is expected to be the case because the spinal columns have been sectioned at varying places within the spinal column for each spinal column and some tissue types are more or less present at different points throughout the spinal column. For example, some areas are more densely packed with nerves than other areas, which results in a higher abundance of ions from that tissue type. These drastic changes, even at small step sizes down the spinal column demonstrate why spatial analysis is important. While slight differences within cell types and states are present, the ions used to discriminate the each spinal cord feature are robust in the fact that even slight differences in depth do not affect their localizations to specific tissue types. For instance, [SM(34:1;O2)+Na]^+^ most notably localizes to the intervertebral disc and blood vessels (Figure 6A), while [PC(O-34:1)+Na]^+^ is primarily present in the bone marrow of two of the spines (Figure 6A, 6B), but it is has a difference in abundance in the white matter of each spine. [PC(O-34:1)+Na]^+^is more abundant in the white matter of the third spine, demonstrating a difference in the relative abundance of, perhaps, a specific cell type (Figure 6C). This lipid was less abundant in the spine shown in figure 6A, and rarely present in the spine in figure 6B. We expect this difference to be due to the sections being taken from different areas in the lumbar region of the three mice. The three spines have a different abundance and spatial location for [SM(34:1;O2)+Na]^+^ because it appears to represent the presence of blood vessels^96^ and intervertebral discs^97^ in the lumbar tissue, which are not consistent in the lumbar region as a function of length^96^. [PC(32:1)+H]^+^ is known to be common in biological tissue^98–101^ and is shown here to be located in the nerves surrounding the white matter in the spinal cord (Figure 5H, 5N, 6). Finally, [PC(32:1)+H]^+^, [PC(O-34:1)+H]^+^, [PC(34:2)+K]^+^, and [PC(38:3)+Na]^+^ remained consistent in spatial location across the three spines. We expect this to be due to each of these lipids having the same function across the three rats and throughout the lumbar region.

**Figure 6.**
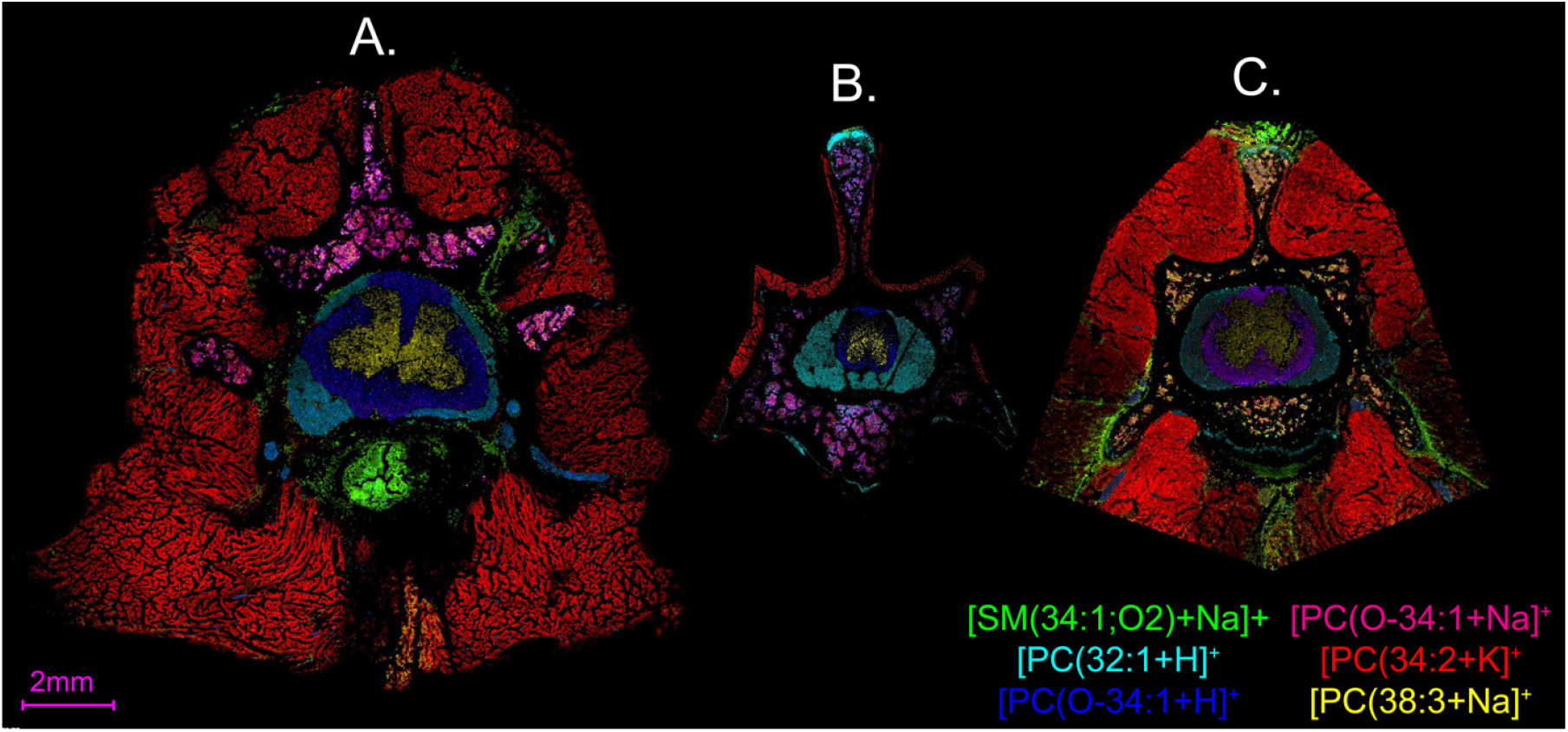
MALDI MSI layered ion images of undecalcified fresh-frozen spinal columns transversely cut from three different adult rats, normalized by total ion count. (A) Spinal column cut transversely with a raster width of 30 μm. (B) Spinal column cut transversely with a raster width of 10 μm and minimal surrounding muscle included in MALDI MSI. (C) Spinal column cut transversely with a raster width of 30 μm.

## CONCLUSION

Using MALDI mass spectrometry imaging (MSI) on undecalcified fresh-frozen spinal column allows us to see the differences in chemical composition and create images of the native spinal column across several tissue types without the assistance of labels, staining, etc. Using H&E staining and MALDI MSI to visualize the resulting sections, we found that our method helps maintain the spatial and structural integrity of the spinal column, ensuring that single-cell data accurately represents the *in vivo* cellular environment. This method can be applied to study genetic diseases and structural defects, by comparing the biomolecules present in disease states versus non-diseased tissue, and observing the biomolecular changes during development or due to traumatic injuries. This method allows for the precise sectioning of undecalcified fresh-frozen spinal columns, providing a valuable tool for studying the spinal column and related diseases. The ability to section undecalcified spinal column should allow for spatial analysis of biomolecules at a single cell level within the spinal column without the interaction or degradation associated with decalcifying bones. Analyzing native bone tissue metabolism at the single-cell level will lead to insights into bone physiology and pathophysiology, as well as potential applications in regenerative medicine and therapeutic development for bone-related disorders.

## ASSOCIATED CONTENT

### Supporting Information

The Supporting Information is available free of charge on the ACS Publications website.

Figure S1 showing additional mass spectra, Figure S2 showing labeled tissue types, Table S1 detailing timsTOF fleX parameters for MALDI MSI, and Table S2 listing lipid assignments (PDF)

## AUTHOR INFORMATION

### AuthorContributions

K.J.B. conducted most of the experiments and performed data analysis. Y.W., Z.S., and A.W. provided the tissue and helped with data interpretation. C.Y.Z. helped with sample preparation. K.J.B. and E.K.N. designed the experiments and participated in data interpretation. E.K.N. and A.W. supervised the research. All authors reviewed the manuscript. The manuscript was written through contributions of all authors.

### Notes

The authors declare no competing financial interest.

## ACKNOWLEDGMENT

The authors would like to thank the University of California, Davis for its financial support. This work was in part supported by NIH grants (5R01NS100761 and 1R01NS115860). Figure 1 was created using BioRender.com (FK25PLSLCF).

## Notes

### Competing Interest Statement

The authors have declared no competing interest.

## REFERENCES

(1) Little, J. P. Chapter 22 - The Spine: Biomechanics and Subject-Specific Finite Element Models. In DHM and Posturography; Scataglini, S., Paul, G., Eds.; Academic Press, 2019; pp 287–293. 10.1016/B978-0-12-816713-7.00022-2.

(2) Kayalioglu, G. The V, ertebral Column and Spinal Meninges. Spinal Cord 2009, 17–36. 10.1016/B978-0-12-374247-6.50007-9.

(3) Heise, C.; Kayalioglu, G. Chapter 13 - Spinal Cord Transmitter Substances. In The Spinal Cord; Watson, C., Paxinos, G., Kayalioglu, G., Eds.; Academic Press: San Diego, 2009; pp 191–208. 10.1016/B978-0-12-374247-6.50017-1.

(4) Brown, A. G. Organization in the Spinal Cord: The Anatomy and Physiology of Identified Neurones; Springer Science & Business Media, 2012.

(5) McDonald, J. W.; Belegu, V.; Becker, D. Chapter 64 - Spinal Cord. In Principles of Tissue Engineering (Fourth Edition); Lanza, R., Langer, R., Vacanti, J., Eds.; Academic Press: Boston, 2014; pp 1353–1373. 10.1016/B978-0-12-398358-9.00064-1.

(6) Mansfield, P. J.; Neumann, D. A. Structure and Function of the Vertebral Column. Essent. Kinesiol. Phys. Ther. Assist. 2019, 178–232. 10.1016/B978-0-323-54498-6.00008-4.

(7) Webborn, N.; Goosey-Tolfrey, V. Chapter 10 - Spinal Cord Injury. In Exercise Physiology in Special Populations; Buckley, J. P., Ed.; Churchill Livingstone: Edinburgh, 2008; pp 309–334. 10.1016/B978-0-443-10343-8.00010-X.

(8) Isaac, Z.; Katz, J. N. Lumbar Spine Disorders. Rheumatol. Sixth Ed. 2015, 1–2, 578–594. 10.1016/B978-0-323-09138-1.00072-3.

(9) Hassan, A.-E. S.; Du, Y. L.; Lee, S. Y.; Wang, A.; Farmer, D. L. Spina Bifida: A Review of the Genetics, Pathophysiology and Emerging Cellular Therapies. J. Dev. Biol. 2022, 10 (2). 10.3390/jdb10020022.

(10) Simonds, A. K. Scoliosis and Kyphoscoliosis. Clin. Respir. Med. Fourth Ed. 2012, 756–762. 10.1016/B978-1-4557-0792-8.00063-5.

(11) Wasserman, B. R.; Moskovicich, R.; Razi, A. E. Rheumatoid Arthritis of the Cervical Spine. Bull NYU Hosp Jt. Dis 2011, 69, 136–148.

(12) Laporte, S.; Van den Abbeele, M.; Rohan, P. Y.; Adam, C.; Rouch, P.; Skalli, W. Spine. Biomech. Living Organs Hyperelastic Const. Laws Finite Elem. Model. 2017, 471–495. 10.1016/B978-0-12-804009-6.00022-5.

(13) Berić, A. Chapter 21 - Spinal Cord Injury. In Handbook of Pain Management; Melzack, R., Wall, P. D., Eds.; Churchill Livingstone: Philadelphia, 2003; pp 329–337. 10.1016/B978-0-443-07201-7.50025-4.

(14) Sloan, T. B. Chapter 183 - Spinal Cord Injury. In Complications in Anesthesia (Second Edition); Atlee, J. L., Ed.; W.B. Saunders: Philadelphia, 2007; pp 737–740. 10.1016/B978-1-4160-2215-2.50188-5.

(15) Ahuja, C. S.; Cadotte, D. W.; Fehlings, M. 33 - Spinal Cord Injury. In Principles of Neurological Surgery (Fourth Edition); Ellenbogen, R. G., Sekhar, L. N., Kitchen, N. D., da Silva, H. B., Eds.; Elsevier: Philadelphia, 2018; pp 518–531.e3. 10.1016/B978-0-323-43140-8.00033-0.

(16) Kim, G. U.; Park, W. T.; Chang, M. C.; Lee, G. W. Diagnostic Technology for Spine Pathology. Asian Spine J. 2022, 16 (5), 764–775. 10.31616/asj.2022.0374.

(17) Choi, B. W.; Choi, M. S.; Chang, H. Radiological Assessment of the Effects of Anterior Cervical Discectomy and Fusion on Distraction of the Posterior Ligamentum Flavum in Patients with Degenerative Cervical Spines. CiOS Clin. Orthop. Surg. 2021, 13 (4), 499–504. 10.4055/cios20262.

(18) Moon, M. S.; Choi, W. R.; Lim, H. G.; Lee, S. Y.; Wi, S. M. Pavlov’s Ratio of the Cervical Spine in a Korean Population: A Comparative Study by Age in Patients with Minor Trauma without Neurologic Symptoms. CiOS Clin. Orthop. Surg. 2021, 13 (1), 71–75. 10.4055/cios19174.

(19) Brouwer, R.; Jakma, T.; Bierma-Zeinstra, S.; Ginai, A.; Verhaar, J. The Whole Leg RadiographStanding versus Supine for Determining Axial Alignment. Acta Orthop. Scand. 2003, 74 (5), 565–568. 10.1080/00016470310017965.

(20) Wang, X.; Chen, X.; Fu, Y.; Chen, X.; Zhang, F.; Cai, H.; Ge, C.; Zhang, W. Analysis of Lumbar Lateral Instability on Upright Left and Right Bending Radiographs in Symptomatic Patients with Degenerative Lumbar Spondylolisthesis. BMC Musculoskelet. Disord. 2022, 23 (1), 59. 10.1186/s12891-022-05017-1.

(21) Leone, A.; Guglielmi, G.; Cassar-Pullicino, V. N.; Bonomo, L. Lumbar Intervertebral Instability: A Review. Radiology 2007, 245 (1), 62–77. 10.1148/radiol.2451051359.

(22) Hounsfield, G. N. Computed Medical Imaging. Science 1980, 210 (4465), 22–28. 10.1126/science.6997993.

(23) Chawla, S. Multidetector Computed Tomography Imaging of the Spine. J. Comput. Assist. Tomogr. 2004, 28.

(24) Blackmore, C. C.; Mann, F. A.; Wilson, A. J. Helical CT in the Primary Trauma Evaluation of the Cervical Spine: An Evidence-Based Approach. Skeletal Radiol. 2000, 29 (11), 632–639. 10.1007/s002560000270.

(25) Holmes, J. F.; Akkinepalli, R. Computed Tomography Versus Plain Radiography to Screen for Cervical Spine Injury: A Meta-Analysis. J. Trauma Acute Care Surg. 2005, 58 (5).

(26) Miglioretti, D. L.; Johnson, E.; Williams, A.; Greenlee, R. T.; Weinmann, S.; Solberg, L. I.; Feigelson, H. S.; Roblin, D.; Flynn, M. J.; Vanneman, N.; Smith-Bindman, R. The Use of Computed Tomography in Pediatrics and the Associated Radiation Exposure and Estimated Cancer Risk. JAMA Pediatr. 2013, 167 (8), 700–707. 10.1001/jamapediatrics.2013.311.

(27) Jandial, R. 74 - Posterior Lumbar Interbody Fusion. In Core Techniques in Operative Neurosurgery (Second Edition); Jandial, R., Ed.; Elsevier: Philadelphia, 2020; pp 403–405. 10.1016/B978-0-323-52381-3.00074-5.

(28) Aniq, H.; Campbell, R. Chapter 13 - Magnetic Resonance Imaging. In Pain Management (Second Edition); Waldman, S. D., Ed.; W.B. Saunders: Philadelphia, 2011; pp 106–116. 10.1016/B978-1-4377-0721-2.00013-1.

(29) Chan, R. W.; Lau, J. Y. C.; Lam, W. W.; Lau, A. Z. Magnetic Resonance Imaging. In Encyclopedia of Biomedical Engineering; Narayan, R., Ed.; Elsevier: Oxford, 2019; pp 574–587. 10.1016/B978-0-12-801238-3.99945-8.

(30) Duran, C.; Sobieszczyk, P. S.; Rybicki, F. J. Chapter 13 - Magnetic Resonance Imaging. In Vascular Medicine: A Companion to Braunwald’s Heart Disease (Second Edition); Creager, M. A., Beckman, J. A., Loscalzo, J., Eds.; W.B. Saunders: Philadelphia, 2013; pp 166–183. 10.1016/B978-1-4377-2930-6.00013-6.

(31) Dodo, K.; Fujita, K.; Sodeoka, M. Raman Spectroscopy for Chemical Biology Research. J. Am. Chem. Soc. 2022, 144 (43), 19651–19667. 10.1021/jacs.2c05359.

(32) Beć, K. B.; Grabska, J.; Huck, C. W. Biomolecular and Bioanalytical Applications of Infrared Spectroscopy – A Review. Anal. Chim. Acta 2020, 1133, 150–177. 10.1016/J.ACA.2020.04.015.

(33) Norris, J. L.; Caprioli, R. M. Analysis of Tissue Specimens by Matrix-Assisted Laser Desorption/Ionization Imaging Mass Spectrometry in Biological and Clinical Research. Chem. Rev. 2013, 113 (4), 2309–2342. 10.1021/CR3004295/ASSET/IMAGES/MEDIUM/CR-2012-004295_0028.GIF.

(34) Gross, J. H. Mass Spectrometry: A Textbook; Springer Science & Business Media, 2006.

(35) Engel, K. M.; Prabutzki, P.; Leopold, J.; Nimptsch, A.; Lemmnitzer, K.; Vos, D. R. N.; Hopf, C.; Schiller, J. A New Update of MALDI-TOF Mass Spectrometry in Lipid Research. Prog. Lipid Res. 2022, 86, 101145. 10.1016/j.plipres.2021.101145.

(36) Drake, R. R.; West, C. A.; Mehta, A. S.; Angel, P. M. MALDI Mass Spectrometry Imaging of N-Linked Glycans in Tissues. In Glycobiophysics; Yamaguchi, Y., Kato, K., Eds.; Springer Singapore: Singapore, 2018; pp 59–76. 10.1007/978-981-13-2158-0_4.

(37) Cillero-Pastor, B.; Heeren, R. M. A. Matrix-Assisted Laser Desorption Ionization Mass Spectrometry Imaging for Peptide and Protein Analyses: A Critical Review of On-Tissue Digestion. J. Proteome Res. 2014, 13 (2), 325–335. 10.1021/pr400743a.

(38) Do, T. D.; Ellis, J. F.; Neumann, E. K.; Comi, T. J.; Tillmaand, E. G.; Lenhart, A. E.; Rubakhin, S. S.; Sweedler, J. V. Optically Guided Single Cell Mass Spectrometry of Rat Dorsal Root Ganglia to Profile Lipids, Peptides and Proteins. ChemPhysChem 2018, 19 (10), 1180–1191.

(39) Caprioli, R. M.; Farmer, T. B.; Gile, J. Molecular Imaging of Biological Samples: Localization of Peptides and Proteins Using MALDI-TOF MS. Anal. Chem. 1997, 69 (23), 4751–4760. 10.1021/ac970888i.

(40) Ryan, D. J.; Spraggins, J. M.; Caprioli, R. M. Protein Identification Strategies in MALDI Imaging Mass Spectrometry: A Brief Review. Curr. Opin. Chem. Biol. 2019, 48, 64–72. 10.1016/J.CBPA.2018.10.023.

(41) Piehowski, P. D.; Zhu, Y.; Bramer, L. M.; Stratton, K. G.; Zhao, R.; Orton, D. J.; Moore, R. J.; Yuan, J.; Mitchell, H. D.; Gao, Y.; Webb-Robertson, B.-J. M.; Dey, S. K.; Kelly, R. T.; Burnum-Johnson, K. E. Automated Mass Spectrometry Imaging of over 2000 Proteins from Tissue Sections at 100-?m Spatial Resolution. Nat. Commun. 2020, 11 (1), 8. 10.1038/s41467-019-13858-z.

(42) Neumann, E. K.; Do, T. D.; Comi, T. J.; Sweedler, J. V. Exploring the Fundamental Structures of Life: Non-Targeted, Chemical Analysis of Single Cells and Subcellular Structures. Angew. Chem. Int. Ed. 2019, 58 (28), 9348–9364. 10.1002/anie.201811951.

(43) Bergman, H.-M.; Lindfors, L.; Palm, F.; Kihlberg, J.; Lanekoff, I. Metabolite Aberrations in Early Diabetes Detected in Rat Kidney Using Mass Spectrometry Imaging. Anal. Bioanal. Chem. 2019, 411 (13), 2809–2816. 10.1007/s00216-019-01721-5.

(44) Taylor, M. J.; Lukowski, J. K.; Anderton, C. R. Spatially Resolved Mass Spectrometry at the Single Cell: Recent Innovations in Proteomics and Metabolomics. J. Am. Soc. Mass Spectrom. 2021, 32 (4), 872–894. 10.1021/jasms.0c00439.

(45) Schubert, K. O.; Weiland, F.; Baune, B. T.; Hoffmann, P. The Use of MALDI-MSI in the Investigation of Psychiatric and Neurodegenerative Disorders: A Review. PROTEOMICS 2016, 16 (11–12), 1747–1758. 10.1002/PMIC.201500460.

(46) Rocha, B.; Cillero-Pastor, B.; Blanco, F. J.; Ruiz-Romero, C. MALDI Mass Spectrometry Imaging in Rheumatic Diseases. Biochim. Biophys. Acta BBA - Proteins Proteomics 2017, 1865 (7), 784–794. 10.1016/J.BBAPAP.2016.10.004.

(47) Chen, Y.; Hu, D.; Zhao, L.; Tang, W.; Li, B. Unraveling Metabolic Alterations in Transgenic Mouse Model of Alzheimer’s Disease Using MALDI MS Imaging with 4-Aminocinnoline-3-Carboxamide Matrix. Anal. Chim. Acta 2022, 1192, 339337. 10.1016/J.ACA.2021.339337.

(48) Qi, Z.; Yang, C.; Liao, X.; Song, Y.; Zhao, L.; Liang, X.; Su, Y.; Chen, Z. F.; Li, R.; Dong, C.; Cai, Z. Taurine Reduction Associated with Heart Dysfunction after Real-World PM2.5 Exposure in Aged Mice. Sci. Total Environ. 2021, 782, 146866. 10.1016/J.SCITOTENV.2021.146866.

(49) Fiorentino, G.; Smith, A.; Nicora, G.; Bellazzi, R.; Magni, F.; Garagna, S.; Zuccotti, M. MALDI Mass Spectrometry Imaging Shows a Gradual Change in the Proteome Landscape during Mouse Ovarian Folliculogenesis. Mol. Hum. Reprod. 2023, 29 (4). 10.1093/MOLEHR/GAAD006.

(50) Mallah, K.; Quanico, J.; Trede, D.; Kobeissy, F.; Zibara, K.; Salzet, M.; Fournier, I. Lipid Changes Associated with Traumatic Brain Injury Revealed by 3D MALDI-MSI. Anal. Chem. 2018, 90 (17), 10568–10576. 10.1021/acs.analchem.8b02682.

(51) Sarkis, G. A.; Mangaonkar, M. D.; Moghieb, A.; Lelling, B.; Guertin, M.; Yadikar, H.; Yang, Z.; Kobeissy, F.; Wang, K. K. W. The Application of Proteomics to Traumatic Brain and Spinal Cord Injuries. Curr. Neurol. Neurosci. Rep. 2017, 17 (3), 23. 10.1007/s11910-017-0736-z.

(52) Rohner, T. C.; Staab, D.; Stoeckli, M. MALDI Mass Spectrometric Imaging of Biological Tissue Sections. Mech. Ageing Dev. 2005, 126 (1), 177–185. 10.1016/J.MAD.2004.09.032.

(53) Nishidate, M.; Hayashi, M.; Aikawa, H.; Tanaka, K.; Nakada, N.; Miura, S. ichi; Ryu, S.; Higashi, T.; Ikarashi, Y.; Fujiwara, Y.; Hamada, A. Applications of MALDI Mass Spectrometry Imaging for Pharmacokinetic Studies during Drug Development. Drug Metab. Pharmacokinet. 2019, 34 (4), 209–216. 10.1016/J.DMPK.2019.04.006.

(54) Schulz, S.; Becker, M.; Groseclose, M. R.; Schadt, S.; Hopf, C. Advanced MALDI Mass Spectrometry Imaging in Pharmaceutical Research and Drug Development. Curr. Opin. Biotechnol. 2019, 55, 51–59. 10.1016/J.COPBIO.2018.08.003.

(55) Aichler, M.; Kunzke, T.; Buck, A.; Sun, N.; Ackermann, M.; Jonigk, D.; Gaumann, A.; Walch, A. Molecular Similarities and Differences from Human Pulmonary Fibrosis and Corresponding Mouse Model: MALDI Imaging Mass Spectrometry in Comparative Medicine. Lab. Investig. 2018 981 2017, 98 (1), 141–149. 10.1038/labinvest.2017.110.

(56) Shariatgorji, M.; Nilsson, A.; Fridjonsdottir, E.; Vallianatou, T.; Källback, P.; Katan, L.; Sävmarker, J.; Mantas, I.; Zhang, X.; Bezard, E.; Svenningsson, P.; Odell, L. R.; Andrén, P. E. Comprehensive Mapping of Neurotransmitter Networks by MALDI–MS Imaging. Nat. Methods 2019, 16 (10), 1021–1028. 10.1038/s41592-019-0551-3.

(57) Girod, M.; Shi, Y.; Cheng, J. X.; Cooks, R. G. Desorption Electrospray Ionization Imaging Mass Spectrometry of Lipids in Rat Spinal Cord. J. Am. Soc. Mass Spectrom. 2010, 21 (7), 1177–1189. 10.1016/J.JASMS.2010.03.028.

(58) Girod, M.; Shi, Y.; Cheng, J. X.; Cooks, R. G. Mapping Lipid Alterations in Traumatically Injured Rat Spinal Cord by Desorption Electrospray Ionization Imaging Mass Spectrometry. Anal. Chem. 2011, 83 (1), 207–215. 10.1021/AC102264Z/SUPPL_FILE/AC102264Z_SI_001.PDF.

(59) Kurosu, K.; Islam, A.; Sato, T.; Kahyo, T.; Banno, T.; Sato, N.; Matsuyama, Y.; Setou, M. Expression and Kinetics of Endogenous Cannabinoids in the Brain and Spinal Cord of a Spare Nerve Injury (SNI) Model of Neuropathic Pain. Cells 2022, 11 (24), 4130. 10.3390/cells11244130.

(60) Monroe, E. B.; Annangudi, S. P.; Hatcher, N. G.; Gutstein, H. B.; Rubakhin, S. S.; Sweedler, J. V. SIMS and MALDI MS Imaging of the Spinal Cord. Proteomics 2008, 8, 3746–3754. 10.1002/pmic.200800127.

(61) Hanrieder, J.; Ewing, A. G. Spatial Elucidation of Spinal Cord Lipid- and Metabolite-Regulations in Amyotrophic Lateral Sclerosis. Sci. Rep. 2014 41 2014, 4 (1), 1–7. 10.1038/srep05266.

(62) Sekera, E. R.; Saraswat, D.; Zemaitis, K. J.; Sim, F. J.; Wood, T. D. MALDI Mass Spectrometry Imaging in a Primary Demyelination Model of Murine Spinal Cord. J. Am. Soc. Mass Spectrom. 2020, 31 (12), 2462–2468. 10.1021/jasms.0c00187.

(63) Goodwin, R. J. A. Sample Preparation for Mass Spectrometry Imaging: Small Mistakes Can Lead to Big Consequences. J. Proteomics 2012, 75 (16), 4893–4911.

(64) Calvano, C. D.; Monopoli, A.; Cataldi, T. R. I.; Palmisano, F. MALDI Matrices for Low Molecular Weight Compounds: An Endless Story? Anal. Bioanal. Chem. 2018, 410, 4015–4038.

(65) Buszewska-Forajta, M.; Rafińska, K.; Buszewski, B. Tissue Sample Preparations for Preclinical Research Determined by Molecular Imaging Mass Spectrometry Using Matrix-Assisted Laser Desorption/Ionization. J. Sep. Sci. 2022, 45 (7), 1345–1361. 10.1002/jssc.202100578.

(66) Sugiura, Y.; Shimma, S.; Setou, M. Thin Sectioning Improves the Peak Intensity and Signal-to-Noise Ratio in Direct Tissue Mass Spectrometry. J. Mass Spectrom. Soc. Jpn. 2006, 54 (2), 45–48. 10.5702/massspec.54.45.

(67) Liu, H.; Zhu, R.; Liu, C.; Ma, R.; Wang, L.; Chen, B.; Li, L.; Niu, J.; Zhao, D.; Mo, F.; Fu, M.; Brömme, D.; Zhang, D.; Gao, S. Evaluation of Decalcification Techniques for Rat Femurs Using HE and Immunohistochemical Staining. BioMed Res. Int. 2017, 2017, 9050754. 10.1155/2017/9050754.

(68) Kawamoto, T.; Kawamoto, K. Preparation of Thin Frozen Sections from Nonfixed and Undecalcified Hard Tissues Using Kawamoto’s Film Method (2020). In Skeletal Development and Repair; Hilton, M. J., Ed.; Methods in Molecular Biology; Springer US: New York, NY, 2021; Vol. 2230, pp 259–281. 10.1007/978-1-0716-1028-2_15.

(69) Seeley, E. H.; Wilson, K. J.; Yankeelov, T. E.; Johnson, R. W.; Gore, J. C.; Caprioli, R. M.; Matrisian, L. M.; Sterling, J. A. Co-Registration of Multi-Modality Imaging Allows for Comprehensive Analysis of Tumor-Induced Bone Disease. Bone 2014, 61, 208–216.

(70) Fujino, Y.; Minamizaki, T.; Yoshioka, H.; Okada, M.; Yoshiko, Y. Imaging and Mapping of Mouse Bone Using MALDI-Imaging Mass Spectrometry. Bone Rep. 2016, 5, 280–285.

(71) Vandenbosch, M.; Nauta, S. P.; Svirkova, A.; Poeze, M.; Heeren, R. M. A.; Siegel, T. P.; Cuypers, E.; Marchetti-Deschmann, M. Sample Preparation of Bone Tissue for MALDI-MSI for Forensic and (Pre) Clinical Applications. Anal. Bioanal. Chem. 2021, 413, 2683–2694.

(72) J. Good, C.; K. Neumann, E.; E. Butrico, C.; E. Cassat, J.; M. Caprioli, R.; M. Spraggins, J. High Spatial Resolution MALDI Imaging Mass Spectrometry of Fresh-Frozen Bone. Anal. Chem. 2022, 94 (7), 3165–3172. 10.1021/acs.analchem.1c04604.

(73) 10x Genomics. H&E Staining & Imaging for Visium Spatial Protocols. https://cdn.10xgenomics.com/image/upload/v1660261285/support-docu-ments/CG000160_DemonstratedProtocol_MethanolFixationandHESt aining_RevC.pdf, 2021.

(74) Sud, M.; Fahy, E.; Cotter, D.; Brown, A.; Dennis, E. A.; Glass, C. K.; Merrill, A. H.; Murphy, R. C.; Raetz, C. R. H.; Russell, D. W.; Subramaniam, S. LMSD: LIPID MAPS structure database. Nucleic Acids Research. 10.1093/nar/gkl838.

(75) Sarmento, M. J.; Llorente, A.; Petan, T.; Khnykin, D.; Popa, I.; Nikolac Perkovic, M.; Konjevod, M.; Jaganjac, M. The Expanding Organelle Lipidomes: Current Knowledge and Challenges. Cell. Mol. Life Sci. 2023, 80 (8), 237. 10.1007/s00018-023-04889-3.

(76) Devaux, P. F. Static and Dynamic Lipid Asymmetry in Cell Membranes. Biochemistry 1991, 30 (5), 1163–1173. 10.1021/bi00219a001.

(77) Neumann, E. K.; Comi, T. J.; Rubakhin, S. S.; Sweedler, J. V. Lipid Heterogeneity between Astrocytes and Neurons Revealed by Single-Cell MALDI-MS Combined with Immunocytochemical Classification. Angew. Chem. 2019, 131 (18), 5971–5975. 10.1002/ange.201812892.

(78) Simons, K.; Vaz, W. L. C. Model Systems, Lipid Rafts, and Cell Membranes. Annu. Rev. Biophys. Biomol. Struct. 2004, 33 (1), 269–295. 10.1146/annurev.biophys.32.110601.141803.

(79) Van Meer, G.; Voelker, D. R.; Feigenson, G. W. Membrane Lipids: Where They Are and How They Behave. Nat. Rev. Mol. Cell Biol. 2008, 9 (2), 112–124. 10.1038/nrm2330.

(80) Janmey, P. A.; Kinnunen, P. K. J. Biophysical Properties of Lipids and Dynamic Membranes. Trends Cell Biol. 2006, 16 (10), 538–546. 10.1016/j.tcb.2006.08.009.

(81) Voet, D.; Voet, J. G.; Pratt, C. W. Principles of Biochemistry, 3. ed., international student version.; Wiley: Hoboken, NJ, 2008.

(82) Miyamoto, S.; Hsu, C.-C.; Hamm, G.; Darshi, M.; Diamond-Stanic, M.; Declèves, A.-E.; Slater, L.; Pennathur, S.; Stauber, J.; Dorrestein, P. C.; Sharma, K. Mass Spectrometry Imaging Reveals Elevated Glomerular ATP/AMP in Diabetes/Obesity and Identifies Sphingomyelin as a Possible Mediator. EBioMedicine 2016, 7, 121–134. 10.1016/j.ebiom.2016.03.033.

(83) Tafesse, F. G.; Ternes, P.; Holthuis, J. C. M. The Multigenic Sphingomyelin Synthase Family. J. Biol. Chem. 2006, 281 (40), 29421–29425. 10.1074/jbc.R600021200.

(84) Taniguchi, M.; Okazaki, T. The Role of Sphingomyelin and Sphingomyelin Synthases in Cell Death, Proliferation and Migration— from Cell and Animal Models to Human Disorders. Biochim. Biophys. Acta BBA - Mol. Cell Biol. Lipids 2014, 1841 (5), 692–703. 10.1016/j.bbalip.2013.12.003.

(85) Mitsutake, S.; Zama, K.; Yokota, H.; Yoshida, T.; Tanaka, M.; Mitsui, M.; Ikawa, M.; Okabe, M.; Tanaka, Y.; Yamashita, T.; Takemoto, H.; Okazaki, T.; Watanabe, K.; Igarashi, Y. Dynamic Modification of Sphingomyelin in Lipid Microdomains Controls Development of Obesity, Fatty Liver, and Type 2 Diabetes. J. Biol. Chem. 2011, 286 (32), 28544–28555. 10.1074/jbc.M111.255646.

(86) Fox, T. E.; Bewley, M. C.; Unrath, K. A.; Pedersen, M. M.; Anderson, R. E.; Jung, D. Y.; Jefferson, L. S.; Kim, J. K.; Bronson, S. K.; Flanagan, J. M.; Kester, M. Circulating Sphingolipid Biomarkers in Models of Type 1 Diabetes. J. Lipid Res. 2011, 52 (3), 509–517. 10.1194/jlr.M010595.

(87) Schlitt, A.; Blankenberg, S.; Yan, D.; Von Gizycki, H.; Buerke, M.; Werdan, K.; Bickel, C.; Lackner, K. J.; Meyer, J.; Rupprecht, H. J.; Jiang, X.-C. Further Evaluation of Plasma Sphingomyelin Levels as a Risk Factor for Coronary Artery Disease. Nutr. Metab. 2006, 3 (1), 5. 10.1186/1743-7075-3-5.

(88) Martínez-Beamonte, R.; Lou-Bonafonte, J.; Martínez-Gracia, M.; Osada, J. Sphingomyelin in High-Density Lipoproteins: Structural Role and Biological Function. Int. J. Mol. Sci. 2013, 14 (4), 7716–7741. 10.3390/ijms14047716.

(89) Signorelli, P.; Conte, C.; Albi, E. The Multiple Roles of Sphingomyelin in Parkinson’s Disease. Biomolecules 2021, 11 (9), 1311. 10.3390/biom11091311.

(90) Kim, M. Y.; Linardic, C.; Obeid, L.; Hannun, Y. Identification of Sphingomyelin Turnover as an Effector Mechanism for the Action of Tumor Necrosis Factor Alpha and Gamma-Interferon. Specific Role in Cell Differentiation. J. Biol. Chem. 1991, 266 (1), 484–489. 10.1016/S0021-9258(18)52461-3.

(91) D’Angelo, G.; Moorthi, S.; Luberto, C. Role and Function of Sphingomyelin Biosynthesis in the Development of Cancer. In Advances in Cancer Research; Elsevier, 2018; Vol. 140, pp 61–96. 10.1016/bs.acr.2018.04.009.

(92) Newton, J.; Lima, S.; Maceyka, M.; Spiegel, S. Revisiting the Sphingolipid Rheostat: Evolving Concepts in Cancer Therapy. Exp. Cell Res. 2015, 333 (2), 195–200. 10.1016/j.yexcr.2015.02.025.

(93) Vanier, M. T. Niemann-Pick Disease Type C. Orphanet J. Rare Dis. 2010, 5 (1), 16. 10.1186/1750-1172-5-16.

(94) Ramakrishanan, N.; Denna, T.; Devaraj, S.; Adams-Huet, B.; Jialal, I. Exploratory Lipidomics in Patients with Nascent Metabolic Syndrome. J. Diabetes Complications 2018, 32 (8), 791–794. 10.1016/j.jdiacomp.2018.05.014.

(95) King, A. M.; Trengove, R. D.; Mullin, L. G.; Rainville, P. D.; Isaac, G.; Plumb, R. S.; Gethings, L. A.; Wilson, I. D. Rapid Profiling Method for the Analysis of Lipids in Human Plasma Using Ion Mobility Enabled-Reversed Phase-Ultra High Performance Liquid Chromatography/Mass Spectrometry. J. Chromatogr. A 2020, 1611, 460597. 10.1016/j.chroma.2019.460597.

(96) Thron, A. K. Vascular Anatomy of the Spine and Spinal Cord. Neurointerventional Manag. Diagn. Treat. 2012, 40.

(97) Agur, A. M. R.; Dalley, A. F. Grant’s Atlas of Anatomy; Lippincott Williams & Wilkins, 2009.

(98) Guo, X.; Cao, W.; Fan, X.; Guo, Z.; Zhang, D.; Zhang, H.; Ma, X.; Dong, J.; Wang, Y.; Zhang, W. Tandem Mass Spectrometry Imaging Enables High Definition for Mapping Lipids in Tissues. Angew. Chem. 2023, 135 (9), e202214804.

(99) Ellis, S. R.; Cappell, J.; Potocnik, N. O.; Balluff, B.; Hamaide, J.; Van der Linden, A.; Heeren, R. M. A. More from Less: High-Throughput Dual Polarity Lipid Imaging of Biological Tissues. Analyst 2016, 141 (12), 3832–3841.

(100) Flinders, B.; Morrell, J.; Marshall, P. S.; Ranshaw, L. E.; Heeren, R. M. A.; Clench, M. R. Monitoring the Three-dimensional Distribution of Endogenous Species in the Lungs by Matrix-assisted Laser Desorption/Ionization Mass Spectrometry Imaging. Rapid Commun. Mass Spectrom. 2021, 35 (1), e8957.

(101) Bowman, A. P.; Heeren, R. M. A.; Ellis, S. R. Advances in Mass Spectrometry Imaging Enabling Observation of Localised Lipid Biochemistry within Tissues. TrAC Trends Anal. Chem. 2019, 120, 115197.

