## Supplemental Information for "Spatial lipidomics of fresh-frozen spines"

*Corresponding Author

### SUPPLEMENTAL INFORMATION

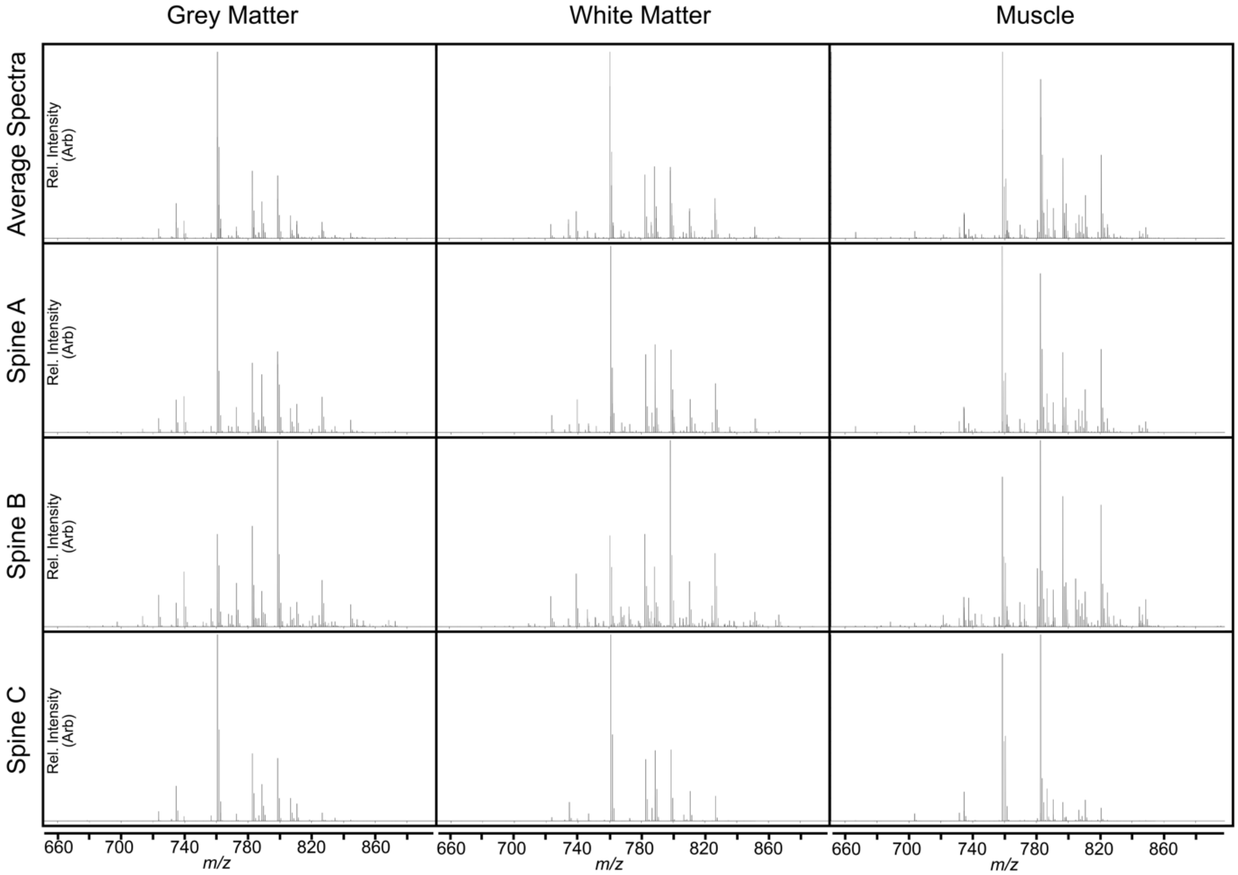


Figure S1. Mass spectra for grey matter, white matter, and muscle tissue in each of three spines and average mass spectra for spines A, B, and C combined for each tissue type.

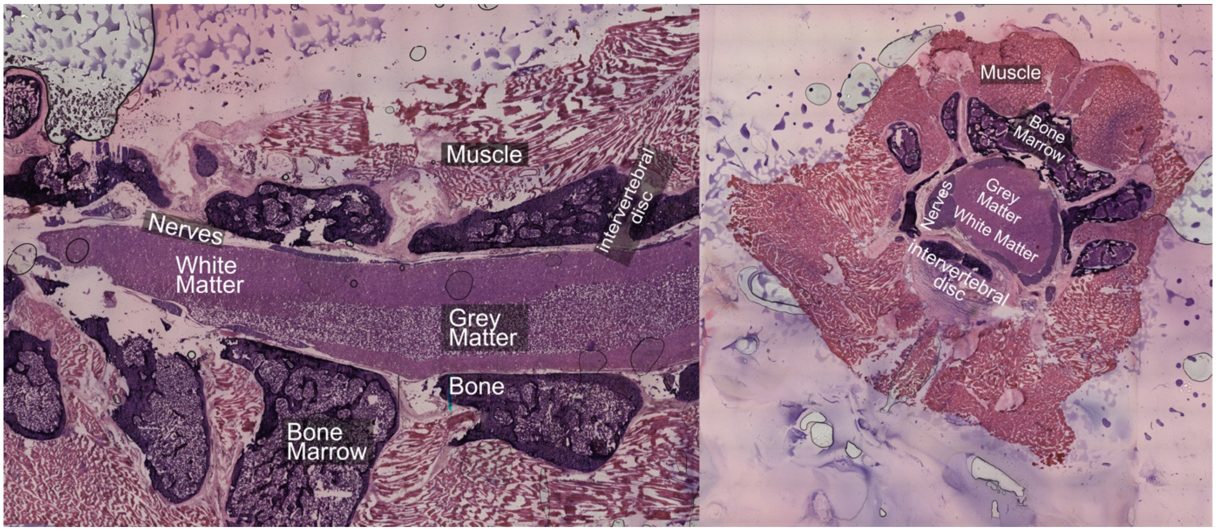

Figure S2. H&E stained sagittal and transverse sections with labels for several tissue types of interest.

| | | |
|---|---|---|
| MS Settings | Scan Begin | 50 *m/z* |
| | Scan End | 1200 *m/z* |
| | Ion Polarity | Positive |
| | Scan Mode | MS |
| Spectrum Settings | Rate Mode | Summation |
| | Rate Value | 165 |
| | MS Averaging | 1 |
| Laser Settings | Laser Power | 88% |
| | Laser Bursts | 1 |
| | Laser shots per burst | 150 |
| | Laser frequency | 10000 Hz |
| Transfer Settings | MALDI Plate Offset | 30.0 V |
| | Deflection 1 Delta | 110.0 V |
| | Funnel 1 RF | 400.0 Vpp |
| | isCID Energy | 0.0 eV |
| | Funnel 2 RF | 400.0 Vpp |
| | Multipole RF | 400.0 Vpp |
| Collision Cell Settings | Collision Energy | 5.0 eV |
| | Collision RF | 500.0 Vpp |
| Quadrupole Settings | Ion Energy | 5.0 eV |
| | Low Mass | 500 *m/z* |
| Focus Pre TOF Settings | Transfer time | 80.0 µs |
| | Pre Pulse Settings | 8.0 µs |
| Detection Settings | High Sensitivity Detection | No |
| | Focus Mode | No |

Table S1. MS settings, spectrum settings, and tune settings for timsTOF fleX.

| Lipid Assignment | *m/z value – Observed* | *m/z value – Calculated* | Error (ppm) |
|---|---|---|---|
| **$[PS(28:1)+H]^+$** | 678.4361 | 678.4341 | 2.9480 |
| **$[SM(34:1;O2)+H]^+$** | 703.5749 | 703.5748 | 0.1421 |
| **$[PC(30:0)+H]^+$** | 706.5384 | 706.5381 | 0.4246 |
| **$[HexCer(32:1;O2)+K]^+$** | 710.4919 | 710.4968 | -6.8966 |
| **$[PA(34:1)+K]^+$** | 713.4501 | 713.4518 | -2.3828 |
| **$[PC(O\text{-}32:0)+H]^+$** | 720.5897 | 720.5902 | -0.6939 |
| **$[PA(36:2)+Na]^+$** | 723.4939 | 723.4935 | 0.5529 |
| **$[SM(34:1;O2)+Na]^+$** | 725.5557 | 725.5568 | -1.5161 |
| **$[SM(36:1;O2)+H]^+$** | 731.6058 | 731.6061 | -0.4101 |
| **$[PC(32:1)+H]^+$** | 732.5535 | 732.5538 | -0.4095 |
| **$[PC(32:0)+H]^+$** | 734.5695 | 734.5694 | 0.1361 |
| **$[PA(36:3)+K]^+$** | 737.4516 | 737.4518 | -0.2712 |
| **$[PA(36:2)+K]^+$** | 739.4673 | 739.4675 | -0.2705 |
| **$[SM(34:1;O2)+K]^+$** | 741.5305 | 741.5307 | -0.2697 |
| **$[PC(O\text{-}34:2)+H]^+$** | 744.5901 | 744.5902 | -0.1343 |
| **$[PC(O\text{-}34:1)+H]^+$** | 746.6041 | 746.6058 | -2.2770 |
| **$[PA(38:2)+Na]^+$** | 751.5228 | 751.5248 | -2.6613 |
| **$[PC(O\text{-}32:2)+K]^+$** | 754.5187 | 754.5147 | 5.3014 |
| **$[PC(32:0)+Na]^+$** | 756.5514 | 756.5514 | 0.0000 |
| **$[PC(34:2)+H]^+$** | 758.5676 | 758.5694 | -2.3729 |
| **$[PC(34:1)+H]^+$** | 760.5859 | 760.5851 | 1.0518 |
| **$[PA(38:2)+K]^+$** | 767.5011 | 767.4988 | 2.9967 |
| **$[PC(O\text{-}34:1)+Na]^+$** | 768.5857 | 768.5878 | -2.7323 |
| **$[SM(36:1;O2)+K]^+$** | 769.5609 | 769.5620 | -1.4294 |
| **$[PC(32:0)+K]^+$** | 772.5248 | 772.5253 | -0.6472 |
| **$[PC(34:2)+Na]^+$** | 780.5513 | 780.5514 | -0.1281 |
| **$[PC(36:4)+H]^+$** | 782.5680 | 782.5694 | -1.7890 |
| **$[PC(34:0)+Na]^+$** | 784.5773 | 784.5827 | -6.8826 |
| **$[PC(36:2)+H]^+$** | 786.5999 | 786.6007 | -1.0170 |
| **$[PC(36:1)+H]^+$** | 788.6163 | 788.6164 | -0.1268 |
| **$[PE(O\text{-}38:5)+K]^+$** | 790.5136 | 790.5147 | -1.3915 |
| **$[PC(O\text{-}36:2)+Na]^+$** | 794.6024 | 794.6034 | -1.2585 |
| **$[PC(34:2)+K]^+$** | 796.5243 | 796.5253 | -1.2555 |
| **$[PC(34:1)+K]^+$** | 798.5427 | 798.5410 | 2.1289 |
| **$[PC(36:4)+Na]^+$** | 804.5500 | 804.5514 | -1.7401 |
| **$[PC(36:3)+Na]^+$** | 806.5675 | 806.5670 | 0.6199 |
| **$[PC(36:2)+Na]^+$** | 808.5815 | 808.5827 | -1.4841 |
| **$[PC(36:1)+Na]^+$** | 810.5983 | 810.5983 | 0.0000 |

| | | | |
|---|---|---|---|
| **$[SM(42:2;O2)+H]^+$** | 813.6830 | 813.6844 | -1.7206 |
| **$[SM(42:1;O2)+H]^+$** | 815.6968 | 815.7000 | -3.9230 |
| **$[PE(44:12)+H-H2O]^+$** | 818.5190 | 818.5119 | 8.6743 |
| **$[PC(36:4)+K]^+$** | 820.5250 | 820.5253 | -0.3656 |
| **$[PC(36:2)+K]^+$** | 824.5566 | 824.5566 | 0.0000 |
| **$[PC(36:1)+K]^+$** | 826.5733 | 826.5723 | 1.2098 |
| **$[PC(40:9)+H]^+$** | 828.5570 | 828.5538 | 3.8622 |
| **$[PC(38:4)+Na]^+$** | 832.5810 | 832.5827 | -2.0418 |
| **$[PC(38:3)+Na]^+$** | 834.5967 | 834.5983 | -1.9171 |
| **$[SM(42:2;O2)+Na]^+$** | 835.6666 | 835.6663 | 0.3590 |
| **$[SM(42:1;O2)+Na]^+$** | 837.6794 | 837.6820 | -3.1038 |
| **$[PC(38:6)+K]^+$** | 844.5260 | 844.5253 | 0.8289 |
| **$[PC(38:5)+K]^+$** | 846.5391 | 846.5410 | -2.2444 |
| **$[PC(38:4)+K]^+$** | 848.5565 | 848.5566 | -0.1178 |
| **$[SM(42:2;O2)+K]^+$** | 851.6475 | 851.6403 | 8.4543 |
| **$[SM(42:1;O2)+K]^+$** | 853.6529 | 853.6559 | -3.5143 |

Table S2. Lipid Assignment with *m/z* values, calibrated quadratically. Calculated using molecular formula of lipid[89].
